# Time in the city: Long-term urban exposure predicts greater exploration and problem-solving in wild red foxes

**DOI:** 10.1101/2025.09.26.678765

**Authors:** F. Blake Morton, Dylan Thompson-Jones, Kristy A. Adaway, Koko M. Sutter, Catia Matos, Georgia B. Freer, Georgia A. A. Fletcher, Carl D. Soulsbury

**Affiliations:** Department of Psychology, University of Hull, Hull, UK; School of Environmental Sciences, University of Hull, Hull, UK; School of Natural Sciences, University of Lincoln, Lincoln, UK

**Keywords:** rapid environmental changes, behavioural innovation, object neophobia, biodiversity crisis, urbanisation, problem-solving ability

## Abstract

Urbanisation is one of the most important forms of human-driven landscape change, altering wildlife populations in unprecedented ways. In terms of behaviour, for example, urbanisation is hypothesised to increase the likelihood of observing urban populations touching, exploring, and solving novel food-related tasks compared to rural areas. However, little is known about the impact of spatiotemporal patterns of urbanisation, particularly historical patterns of change, on these behaviours. We tested this in the world’s most urbanised carnivore, the red fox (*Vulpes vulpes*), by introducing novel food-related tasks (puzzle feeders) to 284 sites throughout Great Britain. We compared tactile and problem-solving behaviours in rural populations, recently colonised urban populations, and long-established urban populations (>40 years). Foxes from 27.4% of locations touched the tasks, foxes from 12.4% of locations solved them. Urban foxes were more likely to touch tasks compared to rural populations. Exploration time, exploratory diversity, and latency to touch tasks did not significantly differ across urban and rural locations. Urbanisation rate from 1994 to 2020 (26 years) did not significantly predict the likelihood of foxes touching or solving tasks across locations. Older urban populations – particularly from London – spent more time exploring tasks and displayed greater exploratory diversity and higher problem-solving success, despite more recent urban populations being equally likely to touch them. Collectively, our findings suggest that certain population characteristics, such as the likelihood of touching/engaging with novelty, potentially emerge early in urbanisation while other characteristics, such as greater exploratory and innovative behaviours, may emerge after long-term urban exposure across many decades.

**Highlights:**

- Historical impacts of urbanisation on wild animal behaviour are unclear.
- We tested this with wild red foxes’ responses to novel food objects.
- Urban foxes were more likely to touch and exploit objects, especially from London.
- Older urban foxes displayed more exploratory and innovative behaviours.
- Length of urban exposure may help predict behavioural responses to novelty.

## Introduction

Earth is presently facing a catastrophic decline in biodiversity, often termed the sixth mass extinction or biodiversity crisis (Ceballos & Ehrlich, 2017; Ceballos et al., 2015). This decline is largely the result of human-driven environmental changes (hereafter “environmental changes”) (Young et al., 2016). One of the most important forms of environmental change on the planet is urbanisation (United Nations Population Division, 2019), defined as a dynamic process characterised by expanding human settlements, such as increased infrastructure (e.g., housing and building densities), human population growth, and reduced habitat heterogeneity (Angel et al., 2011; Grimm et al., 2008). By 2050, over 68% of the world human population is expected to live within cities, compared to only 30% in 1950 (United Nations Population Division, 2019). Such rapid changes expose wildlife to a wide variety of novel situations, such as new or modified habitats (Šálek et al., 2015), anthropogenic resources (Contesse et al., 2004), competitive interactions (Martin & Bonier, 2018), and potential predators and diseases (Guiden et al., 2019; Pedroso-Santos & Costa-Campos, 2020). Most species are experiencing these changes for the first time in their evolutionary history; meaning, evolution has not necessarily equipped them to withstand such unprecedented conditions (Garant, 2020; Gunn et al., 2022; Sih et al., 2011). While many species are at risk of decline or extinction due to slow adaptation, others are unexpectedly thriving for reasons that are still unclear (Finn et al., 2023; Sih et al., 2011). Thus, there is a pressing need to better understand what factors help wildlife respond and adapt to environmental changes, particularly those related to urbanisation.

Several key behaviours have been proposed to help some animals cope with novel situations arising from environmental changes, including bold, exploratory, and innovative behaviour. Bold behaviour can be broadly defined as an animal’s likelihood of engaging with novelty, such as close physical proximity and contact with unfamiliar stimuli, including novel food-related objects (Adaway, Snider, et al., 2025; Bergvall et al., 2011; Breck et al., 2019; Estien et al., 2025; Morton et al., 2023; Takola et al., 2021; Young et al., 2025). Exploration reflects an animal’s tendency to actively investigate a situation, such as the total amount of time or number of behaviours used to physically touch and manipulate novel food-related objects (Adaway, Snider, et al., 2025; Benson-Amram et al., 2013; Daniels et al., 2019; Estien et al., 2025; Morton, 2021; Morton et al., 2021; Sol et al., 2011; Takola et al., 2021). Finally, innovation is commonly defined as using a new or modified behaviour to solve a new or familiar task, such as how successful animals are at extracting food rewards from novel objects (Adaway, Snider, et al., 2025; Benson-Amram et al., 2013; Lee, 1991; Reader & Laland, 2003; Thornton & Samson, 2012).

Greater human-driven environmental changes are believed to increase the likelihood of bold, exploratory, and innovative behaviours being observed within a population (Gunn et al., 2022; Morton et al., 2023; Vincze & Kovacs, 2022) because these behaviors may provide adaptive responses to the novel challenges created by urban expansion (Griffin et al., 2017). Relatively bolder animals, for instance, are more likely to touch and engage with unfamiliar, and therefore potentially risky, resources compared to other organisms (Forss et al., 2015; Griffin et al., 2017; Webster et al., 2009). Animals with greater exploratory and innovative tendencies are often better equipped to gain information and discover solutions to novel challenges (Daniels et al., 2019; Reader & Laland, 2003; Sol et al., 2011; Wild et al., 2019). However, the likelihood of animals displaying such behaviours is not always related to the magnitude of environmental change (Griffin et al., 2017; Gunn et al., 2022; Morton et al., 2023; Vincze & Kovacs, 2022). In terms of urban-rural gradients, for example, many studies find that urban populations are more likely to display bolder and more innovative behaviour than rural populations, while other studies show that rural populations can be more or equally as bold and innovative as their urban counterparts (Breck et al., 2019; Greenberg & Holekamp, 2017; Griffin et al., 2017; Lazzaroni et al., 2024; Morton et al., 2023; Turner et al., 2020; Vincze & Kovacs, 2022). Thus, the relationship between environmental changes and how animals respond to novelty is complex, warranting further research.

One underexplored factor is the long-term historical spatiotemporal patterning of environmental changes, such as regional differences in the rate and length of exposure to changes within landscapes over many decades. In birds, for example, species with longer exposure to urbanisation show reduced fear of people (Mikula et al., 2025). In chimpanzees (*Pan troglodytes*), behavioural diversity and flexibility are predicted by historical patterns of environmental changes from the late Pleistocene to the present day (Kalan et al., 2020). Nevertheless, for the majority of studies, examining the impact of environmental changes on animals’ behavioural responses to novelty has typically been based on contemporary environmental classifications, such as present-day urban-rural gradients (Griffin et al., 2017; Gunn et al., 2022; Morton et al., 2023; Vincze & Kovacs, 2022). While such analyses are important for understanding how different degrees or “magnitudes” of change might impact animal populations (e.g., a steeper gradient suggests greater impact), ecological gradients defined by current environmental conditions lack historical context, which could limit researchers’ ability to predict when, where, and how quickly bold, exploratory, and innovative behaviours are likely to emerge in the future (Kalan et al., 2020; Mikula et al., 2025).

Some studies predict that faster environmental changes impose strong selective pressures favouring rapid behavioural shifts among populations (Griffin et al., 2017; Sih et al., 2011). Thus, populations that are more likely to display certain behaviours as a function of faster historical rates of environmental change (e.g., faster changes in human population, traffic or building density) would support the hypothesis that rapid environmental shifts favour the expression of those behaviours (Sih et al., 2011). By contrast, other studies propose that populations with longer exposure to environmental changes, such as decades-long urban living, may provide animals with the opportunity to adapt incrementally, such as from learning or developmental plasticity (Griffin et al., 2017; Johnson-Ulrich et al., 2021; Sih et al., 2011; Uchida et al., 2016). Thus, populations that are more likely to display certain behaviours as a function of how long they have been exposed to those changes would support the hypothesis that length of exposure is important. Finally, because environmental changes are rarely homogenous and, instead, can vary regionally across different landscapes (United Nations Population Division, 2019), understanding spatial variation in animals’ responses to novelty in relation to historical spatiotemporal changes within the environment could help determine whether behavioural responses are spatially homogenous (e.g., all urban settings) or, instead, more likely to be observed in regions with particular rates or lengths of exposure to those changes, such as particular urban ‘hotspots’.

Red foxes, *Vulpes vulpes*, offer a powerful model for understanding how historical patterns of urbanisation influence present-day behavioural adaptation. As the most globally widespread terrestrial carnivore on the planet (Marsh et al., 2022), foxes were among the first mammals on record to have expanded their ecological niche to urban areas (Harris & Baker, 2001). Reports of urban foxes have existed in British cities at least since the 1930s (Harris & Baker, 2001). In more recent decades, foxes have established themselves in other cities throughout Europe, Asia, Australia, and North America (Contesse et al., 2004; Gil-Fernandez et al., 2020; Plumer et al., 2014; Soulsbury et al., 2010). Although urbanisation is a relatively recent phenomenon in this species’ evolutionary history, notable behavioural and morphological differences are already being documented in urban fox populations compared to the countryside. For instance, urban foxes often have relatively smaller stable isotope dietary repertoires (Scholz et al., 2020), smaller skulls (Parsons et al., 2020), consume anthropogenic food more frequently (Contesse et al., 2004; Fletcher et al., 2025; Scholz et al., 2020), and occupy smaller home ranges (Kobryn et al., 2022; Main et al., 2020) compared to rural fox populations. In terms of bold behaviour, studies find that urban foxes are more likely to consume novel food items (Gil-Fernandez et al., 2020) and physically touch novel food-related objects (Morton et al., 2023) compared to rural foxes. Despite this latter evidence, however, it remains unclear whether urban-rural differences in fox behavioural responses to novelty extend beyond boldness, particularly in terms of their exploration and innovation towards novel food-related opportunities. It is also unknown whether spatial variation in foxes’ behavioural responses to novelty – including bold, exploratory, and innovative behaviours – are reflective of historical spatiotemporal patterns of urbanisation.

Previous work on fox exploration of novel food-related objects found no evidence of urban-rural differences, but was largely confined to just 28 locations primarily from semi-urban and rural areas, which may have limited detectability (Lazzaroni et al., 2024). In terms of fox innovation, previous work has reported no evidence of a significant urban-rural difference, but did not consider the possible role of historical spatiotemporal patterns of urbanisation (Morton et al., 2023). Indeed, in terms of this latter point, the colonisation of urban environments by foxes has been uneven across regions (Scott et al., 2014), and Morton et al. (2023) proposed that foxes inhabiting London – i.e., the oldest urban fox population on record in the UK (Harris & Baker, 2001; Soulsbury et al., 2010) – potentially exhibit heighted likelihoods of bold and innovative behaviour compared to other UK populations. However, spatially explicit geographic analyses, such as Local Indicators of Spatial Association (LISA), are still needed to identify spatial clusters (or “hotspots”) of behaviour to test whether they are predicted by historical patterns of urbanisation.

To address these gaps, we investigated the relationship between historical spatiotemporal patterns of urbanisation and population differences in bold, exploratory, and innovative behaviours among present-day foxes in the UK. We first tested whether present-day urban fox populations differ from rural populations in our target behaviours. Although previous work has already reported similar analyses (Lazzaroni et al., 2024; Morton et al., 2023), our current dataset spans a substantially larger spatial scale, providing an opportunity for a more comprehensive re-evaluation of those prior conclusions. Building on this, we then investigated for the first time three additional questions: First, we tested whether population differences in fox behaviour are predicted by the rate at which urbanisation changed within those areas over the last several decades. Second, we assessed whether foxes’ responses to novelty reflect their length of exposure to urbanisation by comparing the behaviour of populations from long-established urban areas (before 1986) to those from more recently urbanised areas. Finally, using Local Indicators of Spatial Association, we tested whether spatial clusters of fox behaviour exist, and whether those clusters align with the historical rate and duration of urbanisation within those areas. By integrating behavioural data with historical spatiotemporal patterns of urban development, we aim to clarify how past environmental changes potentially shape present-day behavioural strategies, offering new insights into how wildlife responds to urbanisation.

## Methods

### Ethical Note

This research was granted ethical approval by the Animal Welfare Ethics Board of the University of Hull (FHS356) and followed all ASAB/ABS guidelines. Foxes were not handled, trail cameras were placed away from footpaths to reduce public disturbance, and all food items used to attract foxes were safe for ingestion by other species.

### Study Sites and Subjects

We studied 284 locations throughout Great Britain (Figure 1a), including areas within and around many towns and cities (e.g. London, Glasgow, Edinburgh, Bristol, Southampton, and York). Landscapes for these locations were diverse, including public parks, residential gardens, monostand pine/spruce plantations, open fields (including farms), mixed woodland, and coastal and mountainous scrubland. Foxes were not tagged and their participation in our behavioural tests was entirely voluntary. We acquired permission to visit 242 of these locations by contacting city councils and other organizations that owned the land. The remaining 42 locations were accessed by advertising the study through social media and regional wildlife groups; these locations were primarily comprised of golf courses and residential gardens.

**Figure 1.**
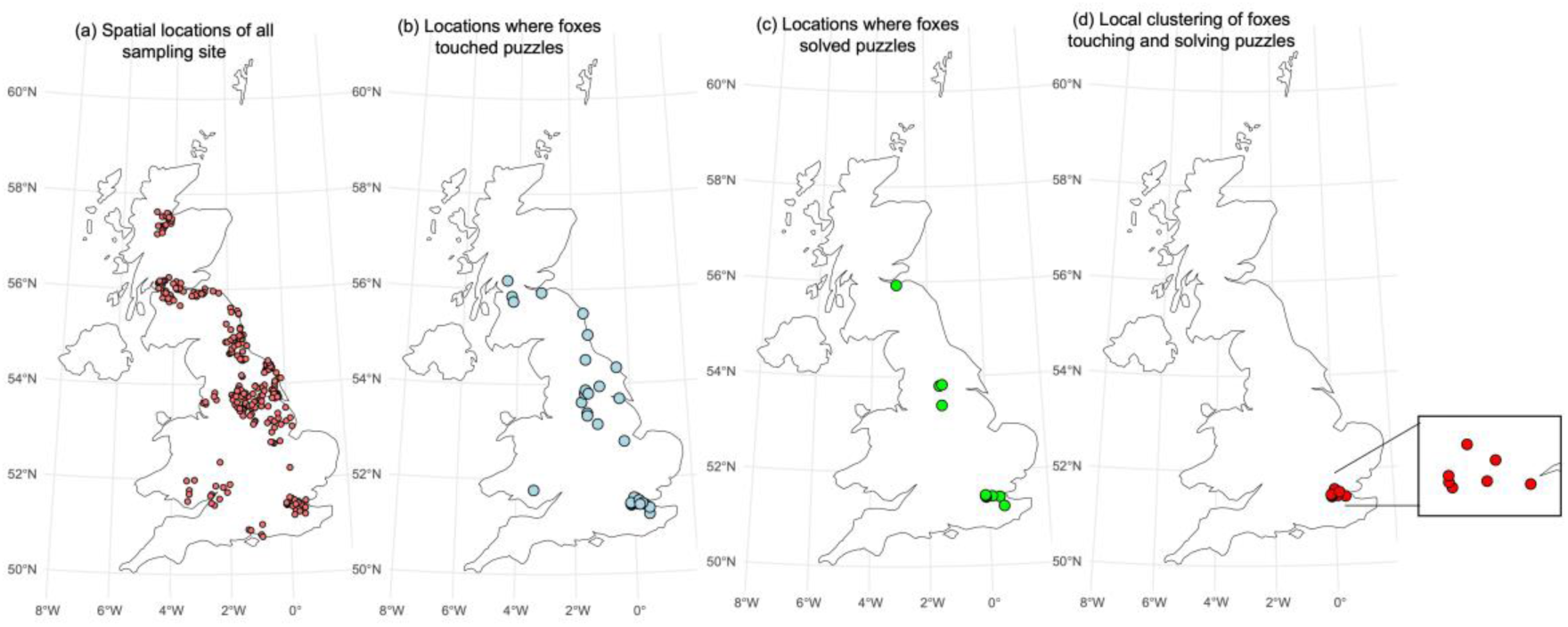
(a) The spatial locations of all sampling sites across the UK (dark red points), (b) the recorded locations where foxes touched the novel tasks (light blue points), (c) the recorded locations where foxes solved the tasks (light green points), and (d) significant (*P*<0.05) local clustering of locations where foxes touched then solved the tasks (red points).

All locations were spaced at least 3.5 km away from any of our other study locations, which reduced the likelihood of acquiring data on the same foxes visiting multiple locations since ≥ 3.5 km is roughly the same or larger than the typical dispersal distance (∼3.7 ± 2.4 km) and home range size (∼1.45 ± 1.5 km^2^) of foxes within the UK (Soulsbury et al., 2011; Trewhella et al., 1988). The same or similar method has been used in other studies of wild animals’ responses to novel objects, such as squirrels, raccoons, and coyotes (Adaway, Snider, et al., 2025; Breck et al., 2019; Chow et al., 2025; Estien et al., 2025; Morton, 2021; Young et al., 2025).

Locations were included in the study based on the following criteria: (1) landowner permission, (2) accessibility to foxes (e.g., no barriers/fences), (3) ability to place our equipment out of public view to avoid theft or vandalism, and (4) the location had to be ≥ 3.5 km from another study area. We had no prior knowledge of fox presence before contacting landowners, and locations were included in the study regardless of whether foxes were known to visit those areas.

### Measuring population-level differences in bold, exploratory, and innovative behaviours in foxes

We used the same methods previously developed to assess population-level differences in foxes’ behavioural responses to novel tasks, the details of which can be found in Morton et al. (2023). As with this other study, we use the term “population-level” in reference to behavioural differences across our urban-rural gradient, where we quantified the likelihood of a behaviour being observed in relation to each location’s specific landscape characteristics.

There were eight novel food-related tasks in total (Figure 2); a single task was selected from the battery and deployed at a given location for 15.6 ± 1.8 days before it was retrieved by researchers. Two weeks is a very typical timeframe for many food-related objects available to British urban foxes, such as regular street cleaning and bin services every 1-2 weeks. Although foxes may respond differently to food-related objects that are left for longer periods, the goal of our study was to measure population differences in *relative* terms, that is, whether urban foxes were more likely to display bold, exploratory, and innovative behaviours compared to rural foxes within the period in which a task was deployed.

**Figure 2.**
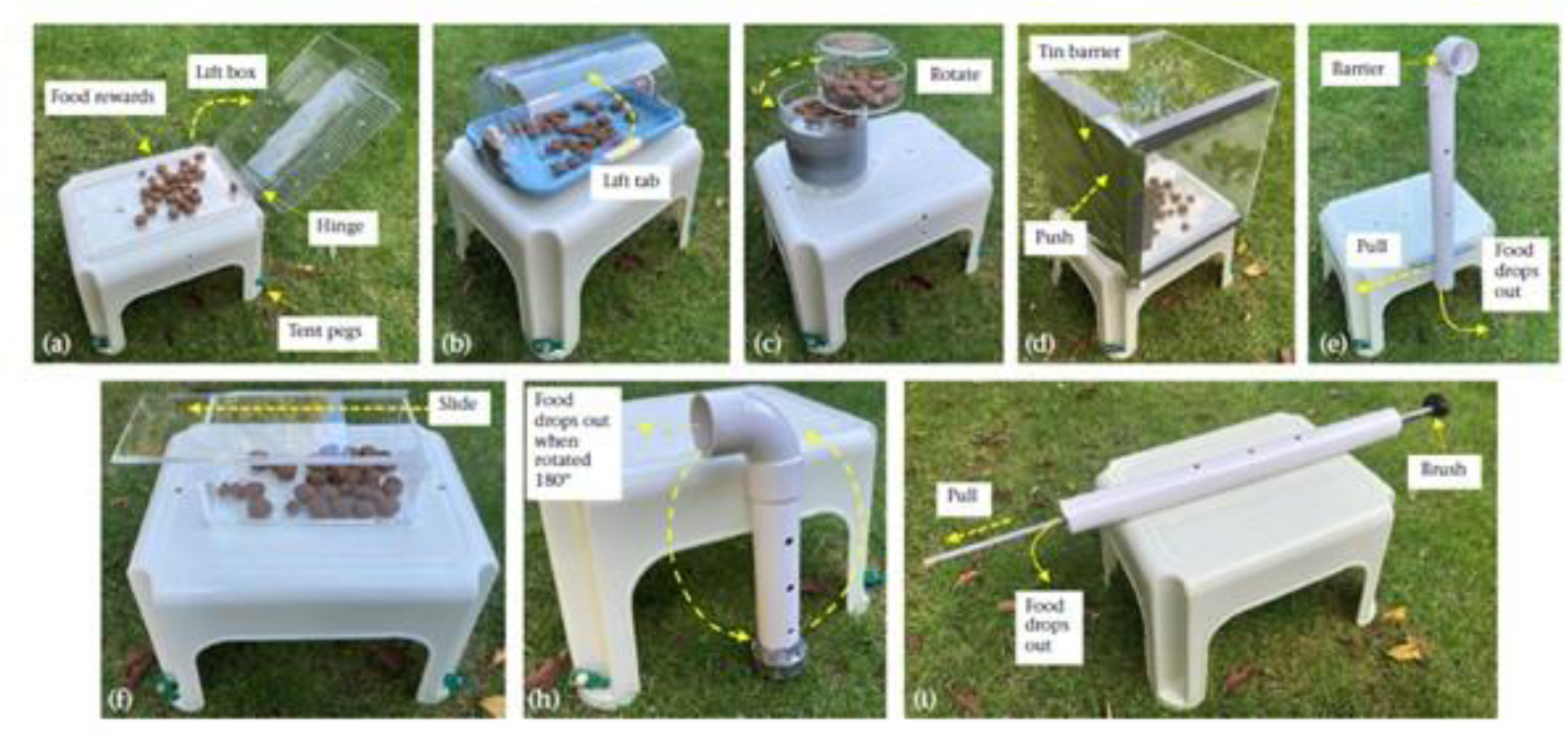
Food-related tasks (objects A, B, C, D, E, F, H and I) presented to foxes. Yellow dashed arrows indicate the direction of each behaviour needed to retrieve the food rewards inside. Object G was never deployed in the field and hence is not depicted in this figure. This figure is from the original Morton et al. (2023) paper.

Data from 200 of our locations come from our original Morton et al. (2023) study, collected between August 2021 and November 2022. Data from an additional 84 locations were collected specifically for the current analysis, between March 2023 and May 2024, and included locations primarily in Wales, Northumberland, Southwest England, and the Scottish Highlands (Figure 1a). Tasks were deployed at up to 25 locations at any given time during 2021-2022 and 2023-2024 due to logistical constraints of obtaining data from 284 locations. With each new set of 25 locations, we counterbalanced a mix of the eight task designs to ensure that our findings were not unique to any particular type of task and, instead, were more generalisable.

Each object varied in terms of design and was made from basic household materials (e.g., PVC piping, metal screws and wooden rods). We also sprayed seventy-five of our tasks with scent deodoriser to understand whether the scent of objects (e.g. human odour) influenced fox behaviour. Although tested in the current study with a larger sample size, our previous work found that, for these specific task designs, population differences in fox behaviour were not significantly influenced by task type or deodoriser (Morton et al., 2023).

All tasks had bait scattered around them to attract foxes to those locations. The objects also contained a food reward inside them, which could be obtained by foxes performing natural behaviours (e.g., lifting, biting, pushing, or pulling materials). Examples of these behaviours can be found here: https://youtu.be/SYHyjXPdcZs (Morton et al., 2023).

Fox visitations occurred when researchers were not present, and ‘no glow’ (940 nm) infrared motion-sensor cameras (Apeman H45) were used to record fox behaviours. Across all locations, we measured whether a fox was detected on camera (1=yes, 0=no) as well as the total number of visits made by foxes during the period in which a task was deployed at that location. We used a 10-minute interval to separate fox detection bouts (hereafter “visits”) following the approach used by Kolowski and Forrester (2017); this provided a practical way to measure the frequency of fox activity at each location. Because foxes were free ranging and their participation was entirely voluntary, we based our analysis on foxes that were at least able and willing to visit the locations, which still enabled us to assess urban-rural differences in foxes’ behavioural responses to the novel tasks (Morton et al., 2023).

We only coded behaviours from locations where foxes clearly acknowledged the task during their visit, which we defined in terms of the fox turning its head to look/smell in the object’s direction. Many factors can potentially underpin whether animals display bold, exploratory, or innovative behaviours (Griffin et al., 2014; Lee & Moura, 2015; Morton, 2021; Reader & Laland, 2003). For example, animals that are too afraid, not hungry, or not persistent may be less likely to exploit novel food-related objects. Importantly, however, the purpose of our study was to determine *whether*, not why, foxes would display our target behaviours in relation to the landscape history of each study location. Thus, as in our previous study (Morton et al., 2023), we measured the likelihood of behavioural innovation occurring within fox populations in terms of the number of locations in which we saw (1=yes, 0=no) a fox displaying a behaviour at any point that led it to operate and successfully gain access to the food inside the novel task within the timeframe in which the task was available at that location. Also, we were not interested in trait boldness in the current study, which describes stable individual differences in behaviour across a wide variety of contexts (Bergvall et al., 2011). Instead, as with Morton et al. (2023) and other studies of free-ranging animals (Adaway, Snider, et al., 2025; Breck et al., 2019; Morton, 2021), we were interested in evaluating the likelihood of fox populations displaying outwardly bold behaviour to exploit unfamiliar food-related objects, which we defined in terms of a fox physically touching (yes or no) the object at any point within the timeframe in which the task was available at that location (e.g., pushing, pulling, licking, or biting it).

Of the foxes that were bold enough to make close physical contact with the novel tasks, we also quantified four additional behavioural measures. First, latency to first touch was defined as the time from the first acknowledgement of a task by any fox at a location until the first instance of physical contact by a fox from that location. Second, to capture foxes’ sustained physical interactions with the task at a given location, we measured the total time foxes spent engaged in tactile exploration of the task, from the very first moment until the very last moment a fox was observed touching the object on all videos. Third, we recorded exploration diversity (Daniels et al., 2019), defined here as the total number of distinct exploratory behaviours directed at the task at any point by foxes during the period in which it was deployed at a location (Table 1). Finally, we measured latency to solve, which we defined in terms of the amount of time elapsed between the first time a fox touched the object until the time when a fox ultimately solved it by gaining access to the food inside, which allowed us to evaluate how long it took a task to be solved, regardless of the total *physical* exploration time.

**Table 1.** Ethogram of behaviours used by foxes to physically explore novel food-related tasks.

| Behaviour | Definition |
| --- | --- |
| Nip | Quick contact with object using teeth, with a fast recoil. Quickly moving body or head away from object after making contact. |
| Bite | Longer contact with teeth compared to a 'nip'. Not moving position of the mouth, staying stationary. Each close of the jaw counts as a new bite. |
| Teeth pull | Grabbing the object with teeth and trying to pull it towards their body. Each close of the jaw counts as new pull. |
| Rostrum touch | Contact with the front of face/nose, which does not include teeth, such as pushing snout into or towards the object, or licking it. |
| Left paw contact | Using left paw to make contact with the object other than standing on the object. |
| Right paw contact | Using right paw to make contact with the object other than standing on the object. |
| Combined paw scratch | Quick contact with the object using both paws at the same time. One paw may make contact first, but both must be touching at some point during the behaviour for combined contact. |

### Recording Fox Behaviour from Trail Cameras

We horizontally placed a ‘no glow’ (940 nm) infrared motion-sensor camera (Apeman H45) approximately 4 m away on a tree trunk at each study location. The cameras had a 120° sensing angle and trigger distance of 20 m. Recording length was set to 5 min (continuous), with a 5 s trigger delay and a 30 s interval between each video. Camera lenses were sprayed with defogger and minor amounts of understory vegetation were removed where necessary to ensure optimal visibility between the camera and task.

### Measuring rates of environmental change due to urbanisation

We obtained spatial data on landscape changes typically associated with urbanisation at each of our study locations from 1994 to 2020, which was the maximum consistent timeframe available. These landscape changes were based on the widely used “LCC: Urban” category from ESA Land Cover Time Series with a resolution of 300 metres (ESA, 2017), which are scores derived from satellite imagery of the presence of built-up areas, including buildings, roads, and other human-made structures. High LCC:Urban scores are also reflective of high human population densities and low cropland densities (Morton et al., 2023). Buffers of 3,500 metres were created around each location to analyse the proportion of urban land cover present; this buffer distance corresponded to the typical annual home range of male and female foxes in the UK (Soulsbury et al., 2011; Trewhella et al., 1988). For each buffer, we calculated the proportion of urban habitat, allowing us to quantify urbanisation in 1994, 2020, and the amount of change between these two time points. Raster data processing was performed with ArcGIS Pro (Version 3.3.1) (Esri Inc., 2024), and landscape cover and change analysis were performed with R version 4.4.2 (Team, 2021).

### Measuring length of exposure to urbanisation

The exact timing of urban foxes colonising different cities is uncertain. We do, however, have a snapshot from a national survey of 378 local UK authorities from 1986 which documented urban fox population status as being either present, absent, or uncertain (Harris & Rayner, 1986). We therefore cross-referenced data on fox presence/absence from our current study with records from 1986; two locations from 1986 were uncertain, both in Hamilton, Scotland, and were excluded, resulting in 54 locations from relatively old urban fox populations and 34 locations from relatively new urban fox populations. Locations where urban foxes were already present in 1986 were defined as relatively “older” urban populations, whereas locations where foxes were absent in 1986 (but present in the current study) were defined as relatively “newer” urban populations.

### Statistical Analyses

All statistical analyses were carried out in R 4.5.0 (R Core Team, 2025). We used Cohen’s kappa tests and intra-class correlation coefficients (ICC 3,1) to evaluate interobserver agreement for all fox behaviours coded from videos. There was excellent agreement across all coders (Tables S1-S8).

We first assessed spatial clustering of foxes touching and then solving the novel tasks using the Local Moran’s I statistic, i.e., a local indicator of spatial association (LISA), using the R package *spdep* (Pebesma & Bivand, 2023). This was based on the locations where foxes were observed touching and solving the tasks, using the behavioural measurements outlined earlier (see ‘*Measuring population-level differences in bold, exploratory, and innovative behaviours in foxes’*). To ensure coverage of the UK, we created a neighbour list using a distance threshold of 35 km to identify spatial relationships between points. The results were visualized using Moran’s scatterplots and maps, highlighting areas of significant spatial autocorrelation, with permutation tests providing p-values to assess the statistical significance of the observed spatial patterns either as areas of high-high (hot spots) or low-low (cold spots) of behaviours.

We structured analyses around *a priori* questions, testing methodological and environmental predictors in separate models. We did not fit a single global model for all of these variables because the ratio of observations to predictors was relatively small. Thus, to avoid model oversaturation, we first ran binomial GLM models to test whether foxes were more likely to engage with or solve the tasks depending on task type, deodoriser spray, camera operation time, and the number of days tasks were deployed (Morton et al., 2023). We also ran a binomial GLM to test whether foxes were more likely to touch or solve the tasks depending on the sequential deployment schedule, that is, whether data were collected during the first (August 2021-November 2022) or the second (March 2023-May 2024) data collection period. Non-significant methodological variables were excluded from our main spatiotemporal analysis to avoid model oversaturation. Finally, we tested whether (a) touching (yes/no) or (b) solving the task (yes/no) was associated with current (2020) urbanisation data using binomial general linear models (GLMs); for both of these variables, we modelled the probability of touching or solving using a spatial generalized linear mixed model (GLMM) with a binomial distribution and logit link.

For behaviours showing significant urban-rural differences, spatial autocorrelation was accounted for by including a Gaussian random field with a Matérn covariance structure (ν fixed at 0.5) using the spaMM package in R (Rousset & Ferdy, 2014). This approach allowed us to test whether any significant urban-rural differences in behaviour were reflective of broadscale urban-rural patterns or, instead, were inflated by local spatial clustering among study locations. Locations’ spatial coordinates were projected to a metric coordinate system (EPSG:27700), and the spatial range and variance were estimated alongside fixed effects (urbanisation).

We used poisson GLMs to examine the effects of urbanisation on the number of visits and exploratory diversity of foxes at each location, followed by gaussian GLMs to assess how urbanisation predicted the time taken to explore or solve the task. Data were log transformed prior to analysis.

We tested whether urbanisation rate was a key driver of behaviour using binomial GLMs, comparing the probability of (a) touching and (b) solving the tasks in relation to the change in LCC:Urban scores at each of our study locations from 1994 to 2020, whereby higher positive values reflect greater rates of change over time. Finally, we compared the likelihood of older versus newer urban fox populations (a) touching and (b) solving the tasks using binomial GLMs, as well as their (c) latency to approach the puzzle (log transformed), (d) time taken to explore puzzle (log transformed), and (e) number of exploratory behaviours using Gaussian and poisson GLMs.

## Results

### Overview of fox behavioural responses to novel tasks

Across all 284 locations, foxes from 46.8% (133/284) locations were detected on cameras. Of these locations, foxes from 93.2% (124/133) locations acknowledged the task. Foxes touched the novel tasks at 27.4% of locations (34/124; Figure 1b) and solved them at 12.4% of locations (14/122; Figure 1c).

The average latency to touch and solve a task was 44.5 ± 71.4 hours and 38.4 ± 71.8 hours, respectively. Total average exploration time was 15.3 ± 20.7 seconds, and the total average number of behaviours used to explore a task was 2.7 ± 1.6 behaviours.

There was no significant effect of task type, deodoriser spray, camera operation time, the number of days tasks were deployed, or the sequential deployment schedule on the likelihood of foxes engaging with or solving the tasks (Tables S9-12). Sample sizes per task were too small to test whether there were differences in time taken to solve the puzzle.

### Behavioural differences between urban and rural fox populations

Number of fox visits to tasks was positively related to urbanisation (LR χ^2^=1205.5, *P*<0.001; Figure S2). For locations where foxes touched a task, there was no effect of urbanisation on the latency to touch (LR χ^2^=0.06, *P*=0.803) or solve (LR χ^2^=0.47, *P*=0.491) the tasks. Urban foxes did not display more exploratory behaviours when interacting with the tasks (LR χ^2^=0.03, *P*=0.866), nor did they spend any longer exploring the tasks (LR χ^2^=0.24, *P*=0.840), compared to rural populations.

### Spatial clustering of fox behaviour

Similar to our previous work (Morton et al., 2023), the likelihood of touching a task was positively associated with urbanisation (Likelihood Ratio (LR) χ^2^=11.05, *P*<0.001; Figure 3a), which remained significant when accounting for spatial autocorrelation (β=16.21 ± 5.28, t=3.07, *P*=0.002). The spatial random field had an estimated range of 1.4 km but negligible variance (λ ≈ 0.29), indicating minimal residual spatial autocorrelation among locations where foxes touched the tasks after accounting for urban cover.

**Figure 3.**
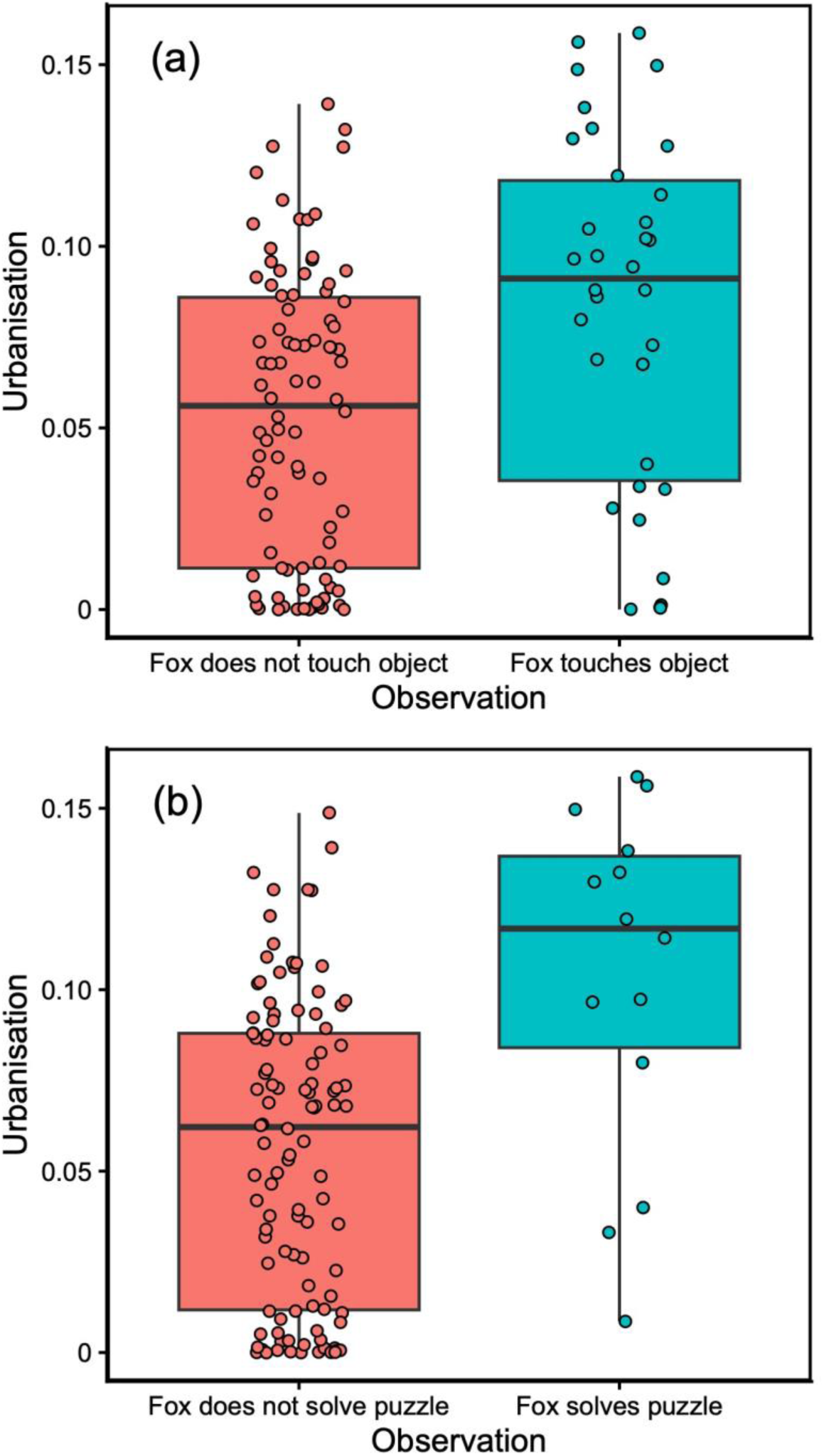
The relationship between urbanisation and whether a fox from a given location (a) does not touch (red) or touches (green) a task, and (b) does not solve (red) or solves (green) a task. The box plots show the median and 25th and 75th percentiles; the whiskers indicate the values within 1.5 times the interquartile range and the circles are data points.

In our extended dataset, there was also a significant positive association between urbanisation and the likelihood of foxes solving a task (LR χ^2^=15.60, *P*<0.001; Figure 3b). This effect, however, was lost when accounting for spatial autocorrelation (β=39.98 ± 114.63, t=0.348, *P*=0.728). The spatial random field had an estimated range of ∼4.8 km and substantial residual variance (λ ≈ 378.0), suggestive of spatial clustering of this behaviour. Foxes mainly solved tasks in urban areas, including Leeds, Edinburgh, Sheffield and London (Figure 1c). The local indicators of spatial association (LISA) identified multiple hotspots for foxes acknowledging the tasks (Figure S1) but only indicated a cluster of 13 significant hot spots for both touching and solving them – all in and around London (Figure 1d).

### Temporal drivers of fox behaviour

The majority of locations (80.2%, 228/284) showed some increase in urbanisation between 1994 and 2020. However, most of these locations had relatively small changes in the amount of urbanisation across this timeframe (mean±SE=0.008±0.004), with only 12 locations showing an increase in urbanisation by >2%. Areas with the greatest increases in urbanisation between 1994 to 2020 did not have a higher likelihood of a fox touching (LR χ^2^=2.82, *P*=0.093) or solving (LR χ^2^=0.01, *P*=0.976) the tasks.

When comparing relatively older versus newer urban fox populations, there was no difference in the likelihood of foxes touching the tasks (LR χ^2^=0.90, *P*=0.341), but older population were significantly more likely to solve them (LR χ^2^=8.11, *P*=0.004). In total, 26.1% (12/46) of foxes from older urban populations solved the tasks compared with 0% (0/16) of foxes from newer populations. Older urban populations did not touch the tasks faster (LR χ^2^=0.20, *P*=0.658) but did spend longer exploring them (LR χ^2^=5.08, *P*=0.024; Figure 4b). Older urban populations also displayed significantly more exploratory behaviours when interacting with tasks (LR χ^2^=9.08, *P*=0.002; Figure 4a). Removing locations in London (identified as a major hotspot) yielded similar results (touch: LR χ^2^=0.27, *P*=0.605; solve: LR χ^2^=4.22, *P*=0.039), indicating that behavioural differences between old versus new urban populations are not solely driven by factors unique to London and, instead, more likely reflect broader patterns of urbanisation among our study sites.

**Figure 4.**
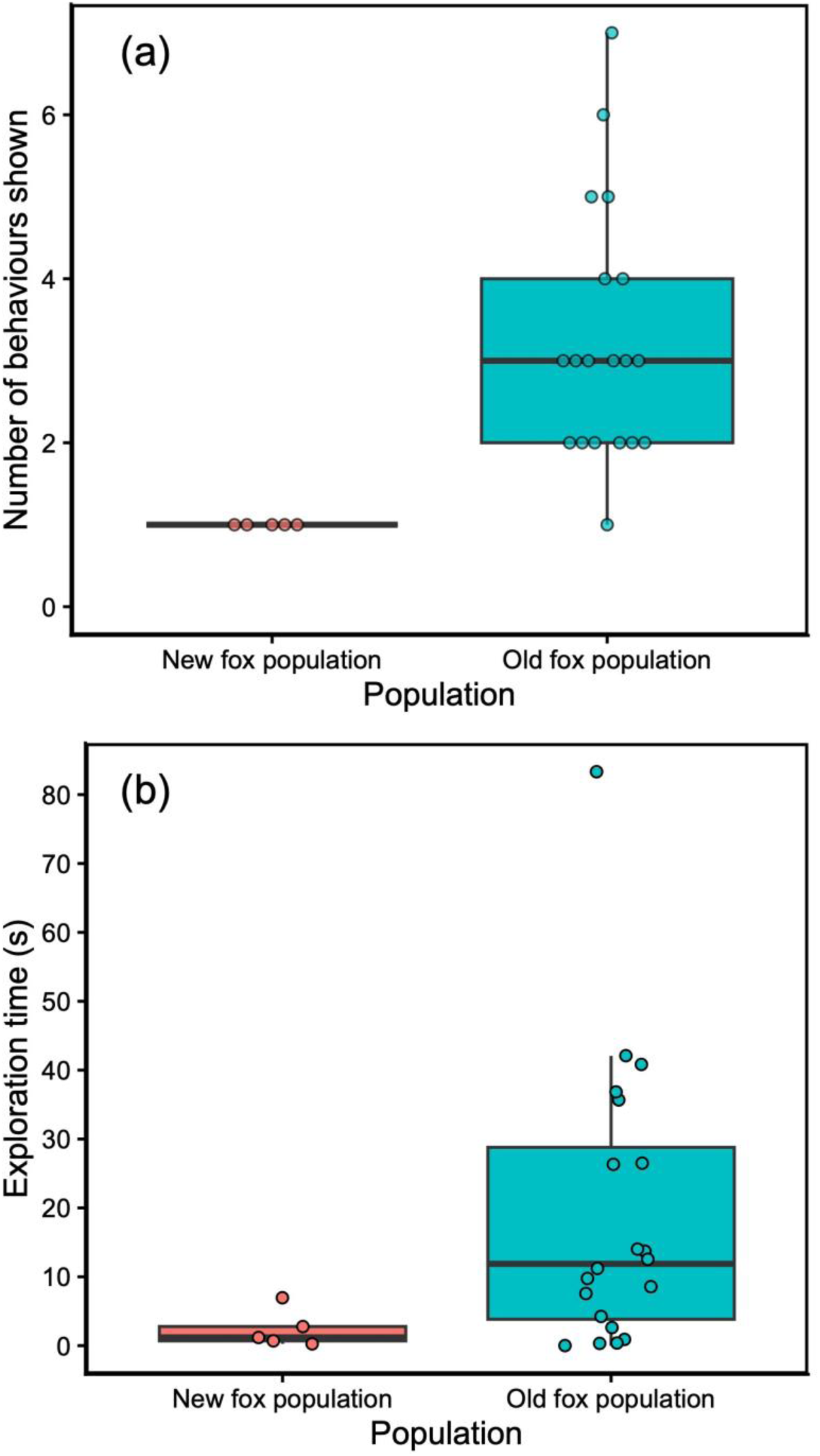
(a) Exploratory diversity (based on the number of unique tactile behaviours from Table 1) displayed by foxes physically interacting with the novel tasks and (b) the time taken to explore the tasks from relatively newer (post-1986) versus older (pre-1986) urban fox populations. The box plots show the median and 25th and 75th percentiles; the whiskers indicate the values within 1.5 times the interquartile range and the circles are data points.

## Discussion

Consistent with our previous findings (Morton et al., 2023), the current study found that urban foxes were more likely to touch novel food-related tasks. We also extend this work by showing, for the first time, that certain urban fox populations are more likely to solve novel tasks using innovative behaviour, and that older, more established urban populations display greater exploration time, exploratory diversity, and problem-solving success compared to more recent urban populations. Collectively, these new findings suggest that historical patterns of urbanisation – specifically, where and how long a city has been inhabited by foxes – may contribute to population differences in bold, exploratory, and innovative behaviours.

Urban foxes were first reported in UK cities in the 1930s (Soulsbury et al., 2010), with 33.1% (125/378) of surveyed urban areas hosting populations by the mid-1980s (Harris & Rayner, 1986). Colonisation has continued, meaning that some of our study populations are older than others (Harris & Rayner, 1986; Wilkinson & Smith, 2001). However, a key finding from the current study is that the rate of urbanisation between 1994 and 2020 did not appear to influence urban-rural differences in our target behaviours, which may reflect the modest changes to urban areas in the UK during this period. Alternatively, this result could reflect the spatial resolution of available landscape datasets, which may not have detected more subtle environmental changes. For example, fine-scale differences in daily human activity patterns, such as traffic or human movement patterns, could influence how fox populations adjust their behaviour in response to those conditions. Regardless, exploratory diversity and problem-solving success were more common in older, more established urban fox populations, even though newer populations (established after 1986 or <40 years ago) were equally likely to touch the tasks. Since foxes first colonised UK cities ∼90 years ago (Soulsbury et al., 2010), these findings suggest that certain behaviours, such as the likelihood of touching/engaging with novelty, may emerge relatively early (<40 years), whereas other behaviours, including greater exploratory and innovative behaviours, may require 40-90 years of urban exposure. In other words, the behavioural characteristics of older urban fox populations suggest a more gradual pattern of adaptation rather than more recent rates of urbanisation, potentially via individual learning, cultural transmission, dispersal patterns, or a combination of these and other mechanisms across many generations. Newer urban populations may not yet exhibit these behaviours to the same extent, being in earlier stages of adjustment or colonisation.

The fact that older urban fox populations tended to be more exploratory and innovative than newer urban populations is inconsistent with the notion that behavioural responses to novelty are particularly advantageous during earlier phases of colonisation (Gruber et al., 2017; Johnson-Ulrich et al., 2021). One possible explanation could be that adaptation to urban environments takes time to emerge in at least some, but not necessarily all, contexts. Indeed, evidence from our study supports this notion. In other species, older urban coot populations (*Fulica atra*) are bolder (Minias et al., 2018) while older commensal house mice populations are better problem-solvers (Vrbanec et al., 2021) compared to younger urban populations. By contrast, Johnson-Ulrich et al. (2021) found that rural hyenas (*Crocuta crocuta*) were more innovative than relatively newer and older urban populations. Whether such differences are explained by genetic or non-genetic factors is not clear (Griffin et al., 2017; Miranda et al., 2013), highlighting the need for longitudinal research to better understand urban-rural divergence in behaviour.

Although urban-rural differences in innovation were observed, these were largely explained by a particular hotspot around London, likely reflecting its status as the oldest urban fox population in the UK (Soulsbury et al., 2010). Once spatial autocorrelation was accounted for, our results aligned with Morton et al. (2023), showing no widespread urban-rural difference. Thus, accounting for regional differences in the length of urban exposure may help explain why urban-rural differences in behaviour are not always consistent between studies (Griffin et al., 2017; Vincze & Kovacs, 2022). Indeed, studies of London foxes have found other distinct phenotypic changes, including skull morphology (Parsons et al., 2020), suggesting that behavioural and morphological phenotypes may be influenced by long-term exposure to urban environments.

Although older urban fox populations were significantly more likely to display more innovative behaviour (even when London was excluded), foxes from Bristol did not currently display innovative behaviours despite being a relatively old urban population (Harris & Rayner, 1986). Historical declines from sarcoptic mange in Bristol during the 1990s (Soulsbury et al., 2007), followed by recolonisation from surrounding rural areas (Baker et al., 2001), may have influenced local factors decreasing foxes’ engagement with novel food-related objects and, hence, the likelihood of innovation. Consistent with this notion, we did not observe bold or exploratory behaviours among current Bristol foxes that were detected on camera. Future research could investigate whether factors related to animals’ local experiences, such as demographic or recolonisation events, potentially “reset” populations to a lower prevalence of bold and exploratory behaviour, affecting the expression of innovative behaviour towards novel food-related objects.

Studies in other species show that bold and innovative behaviours sometimes co-occur within the same population, with novelty avoidance potentially limiting innovation (Mazza & Guenther, 2021; Sol et al., 2011). In our study, innovation required foxes to display sufficient boldness to interact with the tasks, but not all bold fox populations displayed innovative behaviour (e.g., 59% of tasks that were touched were not solved). Because our study focused on population-level behavioural patterns, future research should test whether bold and innovative behaviours co-occur within the same individuals (Mazza & Guenther, 2021).

Urban foxes were more likely to visit our novel tasks, raising the possibility that factors associated with the number of fox visitations, such as differences in opportunities or exposure time to tasks, may have influenced the likelihood of us detecting urban-rural differences in bold and innovative behaviour. However, among locations where foxes interacted with tasks, urban and rural populations did not differ in their latency to touch or solve them, nor did foxes differ in terms of the number of exploratory behaviours or the time spent engaging with tasks, suggesting that number of fox visitations alone cannot fully explain urban-rural differences in bold and innovative behaviour. This notion is further supported by the fact that the overall likelihood of problem-solving was relatively low across all urban and rural fox populations (12.4% of locations), and spatial hotspots for touching and solving were concentrated in specific areas (e.g., London), rather than being more uniform across all high-density urban fox sites (e.g., Glasgow).

Ellington et al. (2024) found that semi-urban vervet monkeys (*Chlorocebus pygerythrus*) selectively explored food-related anthropogenic items more than non-food items. Urban environments likely expose animals to greater availability and diversity of human-made objects, such as discarded litter (DEFRA, 2022), providing more predictable and repeated exposure to these resources (Greggor et al., 2016; Griffin et al., 2017). Such conditions could alter the cost-benefit balance of engaging with unfamiliar human made food-related objects (Lee, 2003), such as the tasks used in our study. Alternatively, innovation may depend on factors beyond what we measured, such as stochastic exploration or the specific type or sequence of exploratory behaviours, which could lead some foxes to solve a task merely by chance. In addition, urban foxes – particularly older established populations – may experience heightened competition over limited resources due to greater population densities compared to rural areas (Soulsbury et al., 2010; Trewhella et al., 1988), which could shape older urban populations’ motivation to explore and exploit novel tasks (Lee & Moura, 2015; Webster et al., 2009).

Our current findings are consistent with other independent data streams, such as human household questionnaires, which report urban-rural differences in fox activity and bold behaviours related to human-fox conflict, and identify London as a key ‘hot spot’ within the UK, including behaviours to exploit humanmade food-related objects (e.g., litter and bin raiding) (Adaway, Hopkins, et al., 2025; Brand & Baldwin, 2020; Dowling, 2013; Harris, 1981; Harris & Baker, 2001; Scott et al., 2014). Nevertheless, because our sample size for fox innovation was relatively small (*N*=34 locations), we recommend future research work towards building larger datasets through collaborative data collection.

## Conclusions

Overall, our study demonstrates that historical patterns of urbanisation, particularly the length of time populations have been established in urban environments, can be an important predictor of population differences in bold, exploratory, and innovative behaviours. While initial engagement with novel food-related opportunities, such as our novel tasks, may emerge relatively quickly in newly established urban fox populations, other behaviours – such as greater problem-solving and length and diversity of exploration – were predominantly observed in older urban populations (particularly London), highlighting the importance of regional differences in the length of exposure to urbanisation in predicting population behaviour. By linking long-term urbanisation history to present-day behavioural traits, our findings provide new insights into the spatial and temporal processes underlying wildlife adaptation to human-altered environments.

## Data Availability

All data are provided in data set S1 in the Supplementary material.

## Declaration of Interest

The authors declare no conflict of interest.

## Acknowledgments

We thank everyone involved with the *British Carnivore Project*, including the researchers, students, and citizen scientists. We are grateful to the many organizations that helped us acquire landowner permissions for the study, particularly staff from the Wildlife Trusts, Scottish Wildlife Trust, The Land Trust, the National Trust, Forestry England (particularly C. Bashforth), Forestry & Land Scotland, Yorkshire Water, and numerous city and regional councils (particularly Hull, East Riding of Yorkshire, and Glasgow). We thank Yacob Haddou (University of Glasgow) for helping us obtain the landscape measures for our study sites. K.A.A. and D.T-J. were both supported at the time of writing by scholarships from the Doctoral College at the University of Hull. K.M.S. was supported at the time of writing by the Leverhulme Doctoral Scholarships Centre for Water Cultures. G.B.F. was supported by a research grant (awarded to C.D.S. and F.B.M.) from the Wild Animal Initiative. F.B.M. also thanks the University of Hull and UKRI Natural Environment Research Council (NERC; Grant No. NE/X018342/1) for funding, and the EU Social Plus programme for providing funding to the University of Hull, which supported three of our research assistants. Finally, we thank the editor and reviewers for their helpful feedback on the manuscript.

## Notes

### Competing Interest Statement

The authors have declared no competing interest.

### Summary of Updates

Clarifications to intro, methods, results, and discussion.

