## Supplementary materials for "Time in the city: Long-term urban exposure predicts greater exploration and problem-solving in wild red foxes"

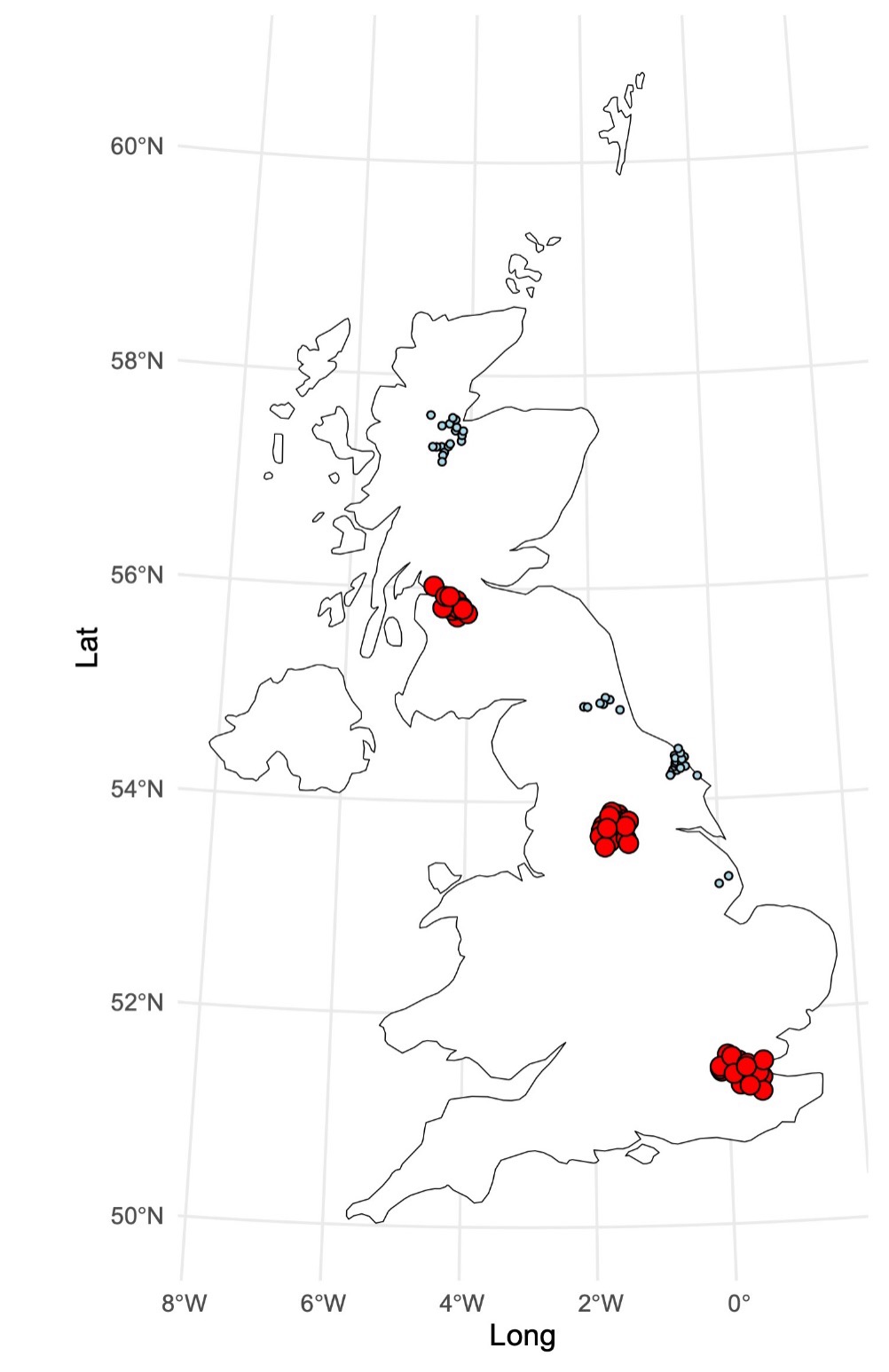

**Figure S1.** The local indicators of spatial association (LISA) multiple hotspots for foxes acknowledging the novel tasks.

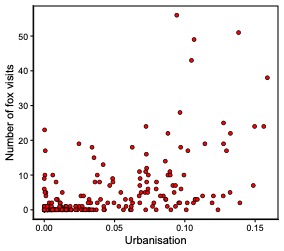

**Figure S2.** The number of visits by foxes to study locations in relation to urbanisation.

**Table S1.** Inter-observer reliability test for foxes detected on camera (1=yes, 0=no).

| Location ID | Coder 1 | Coder 2 |
| --- | --- | --- |
| 222 | 1 | 1 |
| 155 | 1 | 1 |
| 54 | 1 | 1 |
| 3 | 1 | 1 |
| 21 | 1 | 1 |
| 185 | 1 | 1 |
| 159 | 1 | 1 |
| 77 | 1 | 1 |
| 67 | 1 | 1 |
| 12 | 1 | 1 |
| 189 | 1 | 1 |
| 151 | 1 | 1 |
| 143 | 1 | 1 |
| 8 | 1 | 1 |
| 106 | 1 | 1 |
| 112 | 1 | 1 |
| 170 | 1 | 1 |
| 199 | 1 | 1 |
| 125 | 1 | 1 |
| 107 | 1 | 1 |
| 130 | 1 | 1 |
| 10 | 1 | 1 |
| 1 | 1 | 1 |
| 89 | 1 | 1 |

Note. Cohen’s kappa, *k*=1.

**Table S2.** Inter-observer reliability test for fox acknowledgement of novel tasks (1=yes, 0=no).

| Location ID | Coder 1 | Coder 2 |
| --- | --- | --- |
| 222 | 1 | 1 |
| 155 | 1 | 1 |
| 54 | 1 | 1 |
| 3 | 1 | 1 |
| 21 | NA | NA |
| 185 | NA | NA |
| 159 | 1 | 1 |
| 77 | 1 | 1 |
| 67 | 1 | 1 |
| 12 | 1 | 1 |
| 189 | 1 | 1 |
| 151 | 1 | 1 |
| 143 | 1 | 1 |
| 8 | 1 | 1 |
| 106 | NA | NA |
| 112 | 1 | 1 |
| 170 | NA | NA |
| 199 | 1 | 1 |
| 125 | NA | NA |
| 107 | 1 | 1 |
| 130 | 1 | 1 |
| 10 | 1 | 1 |
| 1 | 1 | 1 |
| 89 | 1 | 1 |

Note. Cohen’s kappa, k=1; NA = not applicable (e.g., video was not clear enough to see fox behaviour).

**Table S3.** Inter-observer reliability test of fox touching of novel tasks (1= yes, 0=no).

| Location ID | Coder 1 | Coder 2 |
| --- | --- | --- |
| 222 | 1 | 1 |
| 155 | 0 | 0 |
| 54 | 0 | 0 |
| 3 | 0 | 0 |
| 21 | NA | 0 |
| 185 | NA | NA |
| 159 | 0 | 0 |
| 77 | 0 | 0 |
| 67 | 0 | 0 |
| 12 | 0 | 0 |
| 189 | 0 | 0 |
| 151 | 1 | 1 |
| 143 | 0 | 0 |
| 8 | 0 | 0 |
| 106 | NA | NA |
| 112 | 0 | 0 |
| 170 | NA | NA |
| 199 | 0 | 0 |
| 125 | NA | NA |
| 107 | 0 | 0 |
| 130 | 0 | 0 |
| 10 | 1 | 1 |
| 1 | 0 | 0 |
| 89 | 1 | 1 |

Note. Cohen’s kappa, *k*= 1; NA = not applicable (e.g., video not clear enough to code fox behaviour).

**Table S4.** Inter-observer reliability test for total numbers of unique exploratory behaviours.

| Video | Coder 1 | Coder 2 |
| --- | --- | --- |
| 1 | 1 | 1 |
| 2 | 1 | 2 |
| 3 | 1 | 1 |
| 4 | 1 | 1 |
| 5 | 1 | 1 |
| 6 | 2 | 2 |
| 7 | 3 | 3 |
| 8 | 1 | 1 |
| 9 | 2 | 3 |
| 10 | 4 | 4 |
| 11 | 2 | 3 |
| 12 | 1 | 1 |
| 13 | 3 | 3 |
| 14 | 1 | 1 |
| 15 | 2 | 1 |
| 16 | 1 | 2 |
| 17 | 2 | 3 |
| 18 | 7 | 6 |
| 19 | NA | NA |
| 20 | NA | NA |
| 21 | NA | NA |
| 22 | 1 | 1 |
| 23 | 2 | 2 |
| 24 | 3 | 3 |
| 25 | 3 | 3 |
| 26 | 3 | 1 |
| 27 | 2 | 1 |
| 28 | 1 | 1 |
| 29 | 2 | 2 |
| 30 | 1 | 1 |
| 31 | 1 | 1 |
| 32 | 2 | 2 |
| 33 | 1 | 1 |
| 34 | NA | NA |
| 35 | NA | NA |
| 36 | NA | NA |
| 37 | NA | NA |
| 38 | NA | NA |
| 39 | 2 | 2 |
| 40 | 1 | 1 |
| 41 | 1 | 2 |
| 42 | 2 | 2 |
| 43 | 3 | 3 |

Note. ICC (3,1) = 0.87; NA = not applicable, no exploratory behaviours observed.

**Table S5.** Inter-observer reliability test for the timestamp in which a fox first touched a novel task after it was acknowledged (used to calculate latency to touch).

| Video | Coder 1 | Coder 2 |
| --- | --- | --- |
| 1 | 20/09/2021 (01:01:43) | 20/09/2021 (01:01:41) |
| 2 | 20/09/2021 (01:59:10) | 20/09/2021 (01:59:11) |
| 3 | 20/09/2021 (02:03:24) | 20/09/2021 (02:03:23) |
| 4 | 20/09/2021 (05:01:47) | 20/09/2021 (05:01:46) |
| 5 | 30/09/2021 (04:13:19) | 30/09/2021 (04:13:20) |
| 6 | 16/10/2021 (19:49:39) | 16/10/2021 (19:49:37) |
| 7 | 16/10/2021 (23:48:26) | 16/10/2021 (23:48:36) |
| 8 | 21/01/2022 (07:24:41) | 21/01/2022 (07:24:40) |
| 9 | 23/06/2022 (01:06:04) | 23/06/2022 (01:06:04) |
| 10 | 23/06/2022 (01:31:43) | 23/06/2022 (01:31:43) |
| 11 | 23/06/2022 (03:49:05) | 23/06/2022 (03:49:04) |
| 12 | 24/06/2022 (00:26:28) | 24/06/2022 (00:26:26) |
| 13 | 24/06/2022 (20:50:41) | 24/06/2022 (20:50:36) |
| 14 | 25/06/2022 (23:13:00) | 25/06/2022 (23:12:59) |
| 15 | 26/06/2022 (23:37:23) | 26/06/2022 (23:37:22) |
| 16 | 26/06/2022 (23:40:14) | 26/06/2022 (23:39:32) |
| 17 | 27/06/2022 (23:24:00) | 27/06/2022 (23:23:59) |
| 18 | 30/10/2022 (07:22:02) | 30/10/2022 (07:22:11) |
| 19 | 30/10/2022 (07:28:10) | 30/10/2022 (07:28:08) |
| 20 | 03/11/2022 (03:20:13) | 03/11/2022 (03:20:13) |
| 21 | 08/11/2022 (07:23:16) | 08/11/2022 (07:23:15) |
| 22 | 26/10/2022 (18:41:29) | 26/10/2022 (18:41:32) |
| 23 | 27/10/2022 (06:10:34) | 27/10/2022 (06:10:32) |
| 24 | 28/10/2022 (05:13:32) | 28/10/2022 (05:13:31) |
| 25 | 28/10/2022 (06:51:32) | 28/10/2022 (06:52:01) |
| 26 | 29/10/2022 (18:15:21) | 29/10/2022 (18:15:30) |
| 27 | 31/10/2022 (02:13:52) | 31/10/2022 (02:14:12) |
| 28 | 31/10/2022 (18:33:33) | 31/10/2022 (18:33:31) |
| 29 | 01/11/2022 (00:26:22) | 01/11/2022 (00:26:23) |
| 30 | 07/11/2022 (18:09:33) | 07/11/2022 (18:09:35) |
| 31 | 28/10/2022 (04:18:43) | 28/10/2022 (04:18:42) |
| 32 | 03/11/2022 (21:08:20) | 03/11/2022 (21:08:18) |
| 33 | 29/10/2022 (05:25:29) | 29/10/2022 (05:25:31) |
| 34 | 03/11/2022 (06:34:19) | 03/11/2022 (06:34:10) |
| 35 | 03/11/2022 (06:41:16) | 03/11/2022 (06:41:16) |
| 36 | 05/11/2022 (00:02:20) | 05/11/2022 (00:02:19) |
| 37 | 06/11/2022 (01:46:39) | 06/11/2022 (01:46:43) |
| 38 | 09/11/2022 (23:30:17) | 09/11/2022 (23:30:16) |
| 39 | 26/03/2023 (21:57:37) | 26/03/2023 (21:57:43) |
| 40 | 26/03/2023 (22:04:36) | 26/03/2023 (22:05:40) |
| 41 | 26/03/2023 (22:07:33) | 26/03/2023 (22:07:40) |
| 42 | 02/04/2023 (01:17:51) | 02/04/2023 (01:17:51) |
| 43 | 02/04/2023 (01:22:27) | 02/04/2023 (01:22:35) |

Note. ICC (3,1) = 1.0

**Table S6.** Inter-observer reliability test for the timestamp in which a fox first solved a novel task after it was touched (used to calculate latency to solve).

| Location ID | Coder 1 | Coder 2 |
| --- | --- | --- |
| 10 | 23/08/2021 (05:10:38) | 23/08/2021 (05:10:38) |
| 49 | 02/10/2021 (22:39:25) | 02/10/2021 (22:39:25) |
| 78 | 21/11/2021 (19:10:36) | 21/11/2021 (19:10:36) |
| 138 | 18/02/2022 (21:03:22) | 18/02/2022 (21:03:22) |
| 188 | 23/06/2022 (04:55:51) | 23/06/2022 (04:55:51) |
| 207 | 04/11/2022 (07:02:04) | 04/11/2022 (07:02:14) |
| 209 | 30/10/2022 (07:24:59) | 30/10/2022 (07:24:58) |
| 215 | 04/11/2022 (07:53:57) | 04/11/2022 (07:53:59) |
| 216 | 28/10/2022 (18:22:36) | 28/10/2022 (18:22:34) |
| 218 | 01/11/2022 (02:08:59) | 01/11/2022 (02:08:59) |
| 219 | 28/10/2022 (00:27:21) | 01/11/2022 (05:32:56) |
| 221 | 29/10/2022 (05:25:35) | 29/10/2022 (05:25:37) |
| 226 | 27/03/2023 (13:43:53) | 27/03/2023 (13:43:52) |
| 266 | N/A | N/A |
| 314 | 02/05/2024 (02:36:45) | 02/05/2024 (02:36:48) |

Note. ICC (3,1) = 0.983. NA = not applicable because fox did not solve task.

**Table S7.** Inter-observer reliability test for fox exploration times (seconds).

| Video | Coder 1 | Coder 2 |
| --- | --- | --- |
| 1 | 0.15 | 0.43 |
| 2 | 1.91 | 8.8 |
| 3 | 2.3 | 2.8 |
| 4 | 1.24 | 0.97 |
| 5 | 1.37 | 0.4 |
| 6 | 4.8 | 6.8 |
| 7 | 21.51 | 25.43 |
| 8 | 1.2 | 1.2 |
| 9 | 24.06 | 24.47 |
| 10 | 10.85 | 15.36 |
| 11 | 10.34 | 12 |
| 12 | 1.34 | 1.9 |
| 13 | 4.48 | 7.06 |
| 14 | 0.99 | 1.13 |
| 15 | 2.48 | 3.24 |
| 16 | 0.21 | 1.07 |
| 17 | 15.36 | 16.27 |
| 18 | 35.68 | 38.09 |
| 19 | 0 | 0 |
| 20 | 0 | 0 |
| 21 | 0 | 0 |
| 22 | 1.12 | 0.67 |
| 23 | 2.87 | 3.73 |
| 24 | 18.36 | 8.73 |
| 25 | 8.54 | 3.37 |
| 26 | 3.18 | 0.5 |
| 27 | 0.86 | 0.14 |
| 28 | 1.18 | 1.47 |
| 29 | 20.92 | 2.67 |
| 30 | 26.29 | 23.03 |
| 31 | 0.15 | 0.43 |
| 32 | 0.77 | 1.7 |
| 33 | 0.38 | 1.13 |
| 34 | 0 | 0 |
| 35 | 0 | 0 |
| 36 | 0 | 0 |
| 37 | 0 | 0 |
| 38 | 0 | 0 |
| 39 | 8.27 | 10.12 |
| 40 | 16.39 | 18.67 |
| 41 | 5.42 | 2.31 |
| 42 | 1.39 | 0.57 |
| 43 | 10.61 | 9.35 |

Note. ICC (3,1) = 0.91.

**Table S8.** Inter-observer reliability test for the number of fox visits per location.

| Location ID | Coder 1 | Coder 2 |
| --- | --- | --- |
| 2 | 2 | 2 |
| 4 | 9 | 9 |
| 5 | 13 | 12 |
| 8 | 9 | 8 |
| 12 | 2 | 2 |
| 16 | 0 | 0 |
| 25 | 4 | 4 |
| 34 | 4 | 4 |
| 43 | 1 | 1 |
| 44 | 5 | 5 |
| 54 | 1 | 1 |
| 67 | 1 | 1 |
| 77 | 3 | 3 |
| 82 | 3 | 3 |
| 91 | 17 | 16 |
| 107 | 5 | 5 |
| 130 | 2 | 2 |
| 151 | 7 | 7 |
| 154 | 18 | 18 |
| 156 | 8 | 7 |
| 178 | 18 | 19 |
| 188 | 8 | 8 |
| 198 | 3 | 3 |
| 206 | 15 | 15 |
| 209 | 3 | 3 |
| 220 | 4 | 4 |
| 222 | 5 | 5 |
| 225 | 6 | 6 |
| 259 | 1 | 1 |
| 266 | 10 | 10 |
| 279 | 1 | 1 |
| 297 | 1 | 1 |
| 306 | 2 | 2 |
| 316 | 3 | 3 |

Note. ICC (3,1) = 0.997.

**Table S9.** Binomial GLM analysis between (a) touching and (b) solving the task, and the duration the camera was operational following the fox first acknowledging the task.

| Model | LR 𝛘^2^ | *P* |
| --- | --- | --- |
| (a) Touching | 0.154 | 0.695 |
| (b) Solving | 0.013 | 0.910 |

**Table S10.** Binomial GLM analysis between (a) touching and (b) solving the task, whether data were collected during the first (August 2021-November 2022) or the second (March 2023-May 2024) data collection period.

| Model | LR 𝛘^2^ | *P* |
| --- | --- | --- |
| (a) Touching | 0.898 | 0.343 |
| (b) Solving | 0.738 | 0.390 |

**Table S11.** Binomial GLM analysis between (a) touching and (b) solving the task and whether the number of days the novel tasks were deployed or the presence of a deodoriser impacted the likelihood to touch or solve the tasks.

| Model | Parameter | LR 𝛘^2^ | *P* |
| --- | --- | --- | --- |
| (a) Touch | Deployment time | 1.29 | 0.255 |
|  | Deodoriser | 0.19 | 0.661 |
| (b) Solving | Deployment time | 1.81 | 0.178 |
|  | Deodoriser | 0.77 | 0.379 |

**Table S12.** Binomial GLM models between the type of task deployed at a location and the likelihood of foxes (a) touching, (b) exploring, and (c) solving them.

| Model | LR 𝛘^2^ | *P* |
| --- | --- | --- |
| (a) Touch | 8.77 | 0.269 |
| (b) Exploring | 7.38 | 0.39 |
| (c) Solving | 10.61 | 0.156 |
